# A versatile mouse model to advance human microglia transplantation research in neurodegenerative diseases

**DOI:** 10.1101/2024.11.25.624854

**Authors:** Lutgarde Serneels, Annerieke Sierksma, Emanuela Pasciuto, Ivana Geric, Arya Nair, Anna Martinez-Muriana, An Snellinx, Bart De Strooper

**Author notes:** Equal contributions (Lutgarde Serneels, Annerieke Sierksma, Emanuela Pasciuto, Ivana Geric).

## Abstract

**Background:** Recent studies highlight the critical role of microglia in neurodegenerative disorders, and emphasize the need for humanized models to accurately study microglial responses. Human-mouse microglia xenotransplantation models are a valuable platform for functional studies and for testing therapeutic approaches, yet currently those models are only available for academic research. This hampers their implementation for the development and testing of medication that targets human microglia.

**Methods:** We developed the *hCSF1^Bdes^* mouse line suitable as new transplantation model available to be crossed to any disease model of interest. The *hCSF1^Bdes^* model created by CRISPR gene editing is RAG2 deficient and expresses human CSF1. Additional we crossed this model with two humanized *App* KI mice, the *App^Hu^* and the *App^SAA^ . Flow cytometry, immunohistochemistry and bulk sequencing was used to study the response of microglia in the context of Alzheimer’s disease*.

**Results:** Our results demonstrate the successful transplantation of iPSC-derived human microglia into the brains of hCSF1^Bdes^ mice without triggering a NK-driven immune response. Furthermore we confirmed the multipronged response of microglia in the context of Alzheimer’s disease. The *hCSF1^Bdes^* and the crosses with the Alzheimer’s disease knock-in model *App^SAA^* and the humanized *App knock-in* control mice, *App^Hu^* are deposited with EMMA and fully accessible to the research community.

**Conclusion:** the hCSF1^Bdes^ mouse is available for both non-profit and for-profit organisations, facilitating the use of the xenotransplantation paradigm for human microglia to study complex human disease.

## Background

Roughly 5-10% of our brain cells are microglia (Mittelbronn et al., 2001), yet despite their relatively small number, they are indispensable for proper brain function. These yolk-sac derived brain-resident macrophage related cells undertake a variety of critical functions, including brain surveillance for invading pathogens, removal of debris such as apoptotic cells, myelin or excess synapses, and modulation of neuronal function and synaptic activity. They also secrete a wide array of pro- and anti-inflammatory chemokines and cytokines, ultimately shaping brain function and behavior (Li & Barres, 2018; Munro et al., 2024; Nimmerjahn et al., 2005; Ueno et al., 2013). Microglia are highly dynamic and can adopt complex and diversified molecular, functional and morphological features when confronted with neuropathology (Felsky et al., 2019; Keren-Shaul et al., 2017; Mancuso et al., 2024; Todd et al., 2021; Tuddenham et al., 2024; van den Bosch et al., 2024). Neuropathological features adopted by microglia modify disease progression and severity. To understand how human microglia (hMG) contribute to the etiopathogenesis of neurological disorders, for instance Alzheimer’s disease (AD), it is critical to use appropriate models to expose microglia to pathophysiological relevant conditions.

Microglia are extremely sensitive to external cues and they adapt their morphology and transcriptome very rapidly to changing environmental conditions (Gosselin et al., 2017). Studies of mouse microglia in the brain have been instrumental to understand the complex cell state changes which characterize their responses to amyloid plaque pathology (Deczkowska et al., 2018; Keren-Shaul et al., 2017; Krasemann et al., 2017; Sala Frigerio et al., 2019; Yin et al., 2023). However, the heterogeneity of human microglia is not well recapitulated in mouse microglia (Geirsdottir et al., 2019) hindering the translatability of these findings in AD mouse models to the human disease situation. One third of putative AD risk genes lack adequate mouse orthologs (Mancuso et al., 2019). The transcriptional response of human microglia to AD pathology appears more elaborate, showing strong cytokine responses and antigen presentation reprogramming (Mancuso et al., 2024). This emphasizes the importance of studying human cells to understand human disease (Abud et al., 2017; Capotondo et al., 2017; McQuade et al., 2018). Despite great strides in *in vitro* methods for studying hMG function (Bohlen et al., 2017; Dolan et al., 2023), *in vitro* hMG show a very different expression profile than primary human microglia, e.g. lacking expression of homeostatic markers (e.g. P2RY12, CSF1R, CX3CR1) while upregulating markers of activation (e.g. APOE, GPNMB) (Mancuso et al., 2019).

To overcome these issues, we and others have developed a xenotransplantation model, where hMG are xenografted into the mouse brain. Over time, those hMG colonize and tile the brain, adopt various morphologies depending on their activation status and show a transcriptional profile that is highly akin to primary hMG from human brain (Mancuso et al., 2019, 2024). This model has thus become an essential and powerful tool to study the behavior and function of human microglia (Fattorelli et al., 2021; Hasselmann et al., 2019; Mancuso et al., 2019, 2024; Xu et al., 2020).

Successful transplantation entails a few challenges, including overcoming immune rejection and ensuring the provision of the correct growth factors to allow efficient and long-term engraftment of human microglia (Abud et al., 2017). These requirements thus necessitate an optimal differentiation protocol to ensure the generation of the correct microglia precursor cells able to engraft the brain (Fattorelli et al., 2021).

A key requisite is the inactivation of a functional adaptive immune response by T and B cells (achieved by a NOD SCID or Rag2^-/-^ genetic background) or innate NK cell activity (achieved through Il2rg^-/-^) (e.g., NOD SCID;Il2rg-/- (NSG) or Rag2^-/-^;Il2rg^-/-^) (Shultz et al., 2007). Additionally, these models are engineered to express human cytokines (e.g. -CSF1 alone or in combination with IL3, CSF2) and thrombopoietin (THPO) supporting the development and function of monocytes, macrophages and NK cells (Abud et al., 2017). Hasselmann and Mancuso demonstrated that the expression of human IL3, CSF2 and THPO were not essential for survival and functional development of human microglia (Hasselmann et al., 2019; Mancuso et al., 2019, 2024). By genetically ablating the mouse microglia, full hMG chimerism can be achieved, but this requires the additional removal of the Fms intronic regulatory element (FIRE-enhancer) within the *Csf1R* locus of mouse genome. (Baligács et al., 2024; Munro et al., 2024; Rojo et al., 2019). The generation of these host mice requires the combination of multiple transgenes and makes it complex and expensive to generate those lines. Furthermore, patent restrictions (Regeneron patent EP2675271B1) on certain gene combinations (e.g. Rag2^-/-^, Il2rg^-/-^) found in the available Rag2^tm1.1Flv^; Csf1^tm1(CSF1)Flv^;Il2rg^tm1.1Flv/J^ mouse make that current models are not freely circulating among the researchers needing the models and hindering industry-academia collaborations for pre-clinical therapy development.

To overcome these issues and enhance future studies involving the xenotransplantation of hMG in mouse brain, we generated a new immune-deficient mouse model named hCSF1^Bdes^, which combines *Rag2^-/-^* and human *CSF1* knock in (KI) on a C57Bl6j background, the most widely used mouse strain for modelling human diseases through genetic modifications. This makes our model particularly attractive for crossbreeding with various human disease models, minimizing genetic background variations that could otherwise influence experimental outcomes. We demonstrate here that the combination with *Il2rg*^-/-^ is dispensable for hMG xenotransplantation. We bred hCSF1^Bdes^ onto 2 widely available models used for AD research, *App^Hu^* and *App^SAA^* (Serneels et al., 2020; Xia et al., 2022) and show that xenografted hMG are able to mount the previously described amyloid response in these new mouse models (Mancuso et al., 2024).

## Material and methods

### Mice

***hCSF1 knock in***: Crispr/Cas9 technology was used to replace the genomic DNA sequence corresponding to amino acid 33-552 of the mouse *Csf1* gene with the human *CSF1* genomic sequence corresponding to amino acid 33-554 in the endogenous gene. The 5’ murine signal peptide (amino acid 1-32) and 3’ region of the mouse were retained. The gRNA’s (target sequence ACAGTGTTCTGACACCTCCTTGG and GTGGAACTGCCAGTATAGAAAGG) to the mouse Csf1 gene, the donor vector (PCR assembled from BAC: RP23-15F10 (Mouse) and BAC: RP11-101M23 (Human)), and Cas9 mRNA were co-injected into fertilized mouse eggs of C57BL6J mice. F0 founder animals were identified by extensive PCR analysis followed by Sanger sequence analysis. F0 were bred to wildtype mice to test germline transmission and generate F1 animals. F1 animals were backcrossed to wildtype mice to generate F2 (Additional Figure 1A). These line is called Csf1^em1(CSF1)Bdes^.

**Figure 1.**
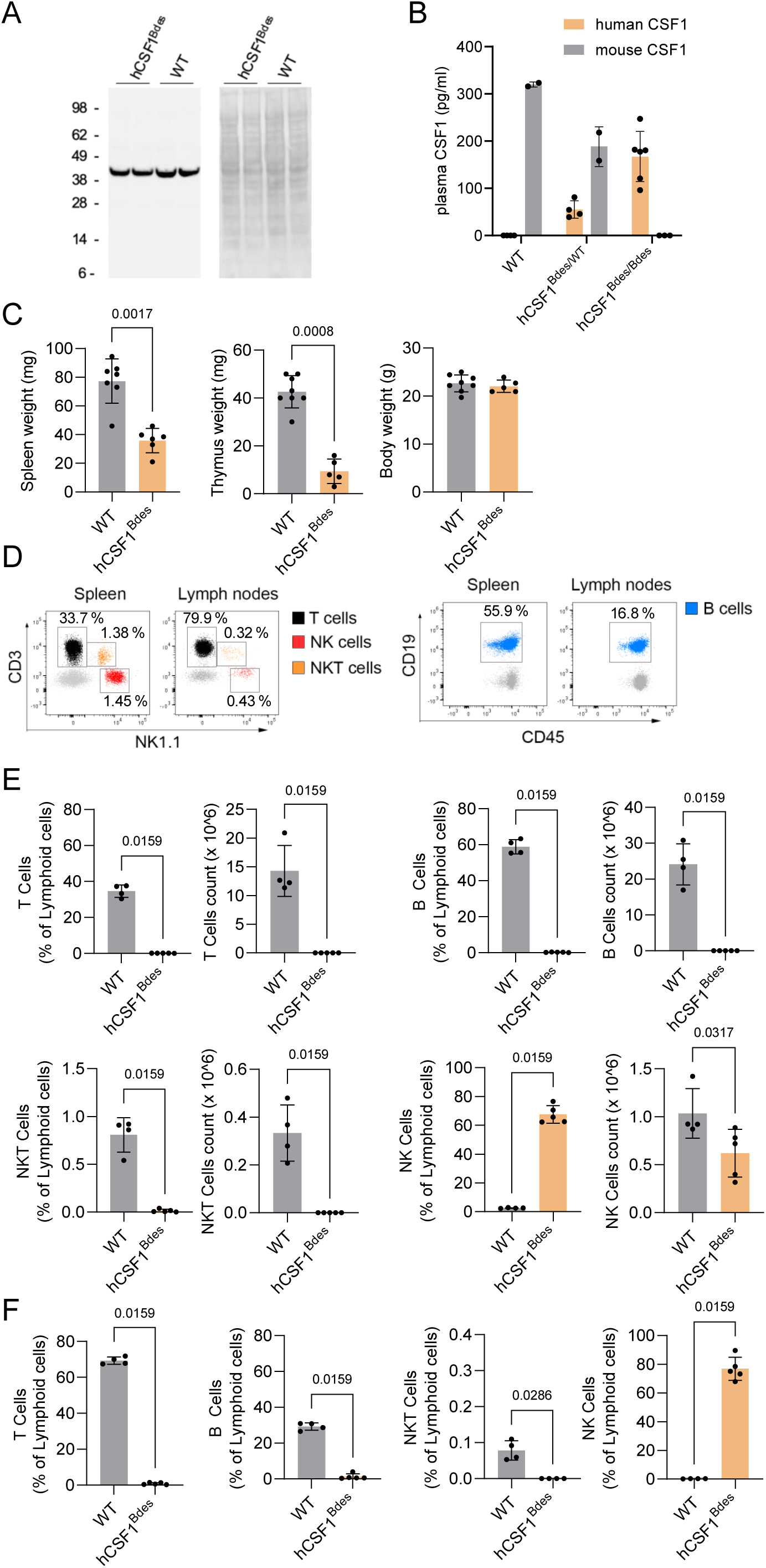
Characterization of CSF1 expression and Rag2-/- effects in hCSF1^Bdes^ mice. A. CSF1 expression in brain of homozygous hCSF1^Bdes^ and WT mice (mixed sexes, 11 weeks of age) was assessed by western blot analysis with an antibody recognizing both human and rodent CSF1 (left panel), showing equal expression levels. The right panel represent the Ponceau S staining as loading control. Position of molecular weight markers are indicated. B. CSF1 expression in the plasma from homozygous and heterozygous CSF1^Bdes^ and WT mice measured with ELISA. Mean, STDEV and N =2-6 are presented on the graph. C. Spleen, thymus and body weight in hCSF1^Bdes^ mice compared to Wild type (WT; C57BL/6j) mice by Mann Whitney test. N=6-7, Bar plots represent mean ± SD. D-F. FACS analysis of lymphocytes and NK cells in hCSF1^Bdes^ mice. (mixed sexes, 11 weeks of age). Single cells were isolated from the spleen and lymph nodes and stained with anti-CD3, anti-CD19, anti-NK1.1, anti-CD45 and anti-CD11b antibodies, and analyzed by flow cytometry. (D) Representative gating strategies to identify T-, B-, NK T cells and NK cells in spleen (left) and Lymph nodes (right) in WT mice. The histograms represent (E) the distribution and total count of lymphoid cells in spleen, and (F) their distribution in lymph nodes. N=6-7, Bar plots represent mean ± SD. hCSF1^Bdes^ mice are compared to the WT mice by Mann Whitney test.

***Rag2 knock out:*** Crisper/Cas9 technology (target sequence AACATAGCCTTAATTCAACCAGG and ACGAGTCTGTAACCGGCTACTGG and Cas9 enzyme) was used to generate an F0 founder in C57Bl6J mice. The F0 founder was selected by extensive analysis utilizing PCR and sequencing which demonstrated a 1198 bp genomic DNA deletion which corresponds to amino acid 16-430 and generates a premature stop (Additional Figure 1B). The F0 founder was backcrossed two times to wildtype mice and the F2 generation was used to generate Csf1^em1(CSF1)Bdes^; Rag2^em1Bdes^ mice, hereafter named Csf1^Bdes^.

Standard genotyping is done by PCR with the following primers: Csf1^Bdes^ allele, AGAGTTAGACAGTTCTGGGTCTTT and ACTGGGTACCTGGAGATAGTT resulting in an amplicon of 252 bp), Csf1^WT^ allele, CCCTTGCCCTAAGGAGATAAA and ACTGGGTACCTGGAGATAGTT resulting in an 355 bp amplicon, Rag2^Bdes^ allele, AGTGTCTGTGATTTCCTAACGG and ACAATAGATCATGGCGGGTTTA resulting in an amplicon of 320 bp, Rag2^WT^ allele GAGATGTCCCTGAACCCAGA and TTGAGTGAGGATTGCACTGG resulting in an amplicon of 404 bp.

**App knock in** : The new xenograft model mouse was crossed with: (1)APP^em1Bdes^ harbouring G676R, F681Y, R684H amino acid substitutions which humanize the Aβ sequence, previously extensively characterized and proposed here as control model (Serneels et al., 2020). We call this cross App^Hu^. (2) With App^tm1.1Dnli^ (Jax strain 034711) harbouring G676R, F681Y, R684H amino acid substitutions to humanize the Aβ sequence and the KM670/671NL (Swedish), E693G (Arctic), and T714I (Austrian) amino acid substitutions which are well known FAD causing mutations (Xia et al., 2022). These mice are called App^SAA^. All the mouse lines and crosses are deposited with EMMA and free to use.

Rag2tm^1.1Flv^; Csf1^tm1(CSF1)Flv^;Il2rg^tm1.1Flv/J^ (Jax strain 017708), named hCSF1^Flv^, and Rag2^tm1.1Cgn^ (Jax strain 008449), named Rag2^Cgn^ and C57BL6JRj (Janvier labs), named WT, were used during the study in control experiments. Both male and female mice have been included in this study. Mice were analysed at the indicated time (see Figure legends). Mice had access to food and water ad libitum and were housed in an SOPF+ facility with a 14/10-h light/dark cycle at 21 °C and 32% humidity, in groups of 2–5 animals. All experiments were conducted according to protocols approved by the local Ethical Committee of Laboratory Animals of KU Leuven (P125-2022).

### Cell culture

Human embryonic stem cells (hESC; WA09 WiCell Research Institute, CVCL 9773) and human induced pluripotent stem cells (iPSCs; UKBi011-A-3, EBiSC) were differentiated into microglia progenitors according to the MIGRATE protocol (Fattorelli et al., 2021). After culturing the stem cells to 70-80% confluency, cells were detached and plated at 10 000 cells/well into a U-bottom 96-well plate. Cells were plated in mTeSR1 media (Stemcell Technologies, 85850) supplemented with 50ng/ml BMP4 (PeproTech, 125-05), 50ng/ml VEGF (PeproTech, 100-20) and 20ng/ml SCF (PeproTech 300-07) to induce embryonic body (EB) formation. On day 4 after plating, EBs were collected and transferred into a 6-well plate in X-VIVO 15 media supplemented with 50 ng/ml SCF (PeproTech 300-07), 50 ng/ml M-CSF (PeproTech, 300-25), 50 ng/ml IL-3 (PeproTech, 200-03), 50 ng/ml Flt3-ligand (PeproTech, 300-19) and 5 ng/ml TPO (PeproTech, 300-18). To induce myeloid lineage producing progenitors, EBs cell culture media was replaced with X-VIVO 15 media supplemented with 50 ng/ml Flt-3-ligand (PeproTech, 300-19), 50 ng/ml M-CSF (PeproTech, 300-25) and 25 ng/ml GM-CSF (PeproTech, 300-03) on day 11 of the protocol. On day 18 of the protocol, progenitors were harvested in PBS and transplanted into mouse brain. Throughout the protocol, cells were maintained in a humidified incubator at 37 °C and 5% CO2.

### Xenotransplantation of human microglia progenitors

To deplete endogenous mouse microglia, newborn mice were injected intraperitoneally with 200mg/kg body weight CSF1R inhibitor BLZ945 (Seleck, S7725) at the age of P2 and P3. At P4, mice were injected with microglia progenitors collected as described above on day 18 according to the MIGRATE protocol (Fattorelli et al., 2021). Mice were first anesthetized by hypothermia, after which they received bilateral injection (coordinates from bregma: anteroposterior, −1 mm; mediolateral, ±1 mm) of cell suspension containing progenitors (1µl, 250 000 cells/µl). After recovering under an infrared lamp, pups were transferred back to their mothers.

### Human microglia isolation

Mice were sacrificed by an overdose of sodium pentobarbital and subsequently perfused with ice-cold heparinized PBS. Brains were dissected into the left and right hemisphere, where one hemisphere was fixed in 4% PFA, and the other was processed for tissue homogenization and hMG isolation. Brain tissue was enzymatically dissociated using the Neuronal Dissociation Kit (P) (Miltenyi, 130-092-628) following manufacturer’s instructions. After dissociation, tissue was passed through a 70um cell strainer and pelleted for 15min at 300g at 4°C. To remove myelin, pellets were resuspended in 30% Percoll solution and centrifuged for 15min at 300g at 4°C, creating a myelin layer, which was carefully removed. The remaining cell pellet was washed with FACS buffer (2% FBS (Life Technologies, 10270106)), 2 mM EDTA (Sigma-Aldrich, E7889) in PBS) and blocked with FcR blocking solution (1:10, Miltenyi, 130-092-575) for 10min at 4°C. As a last step, cells were stained with viability dye eFluor780 (1:1000, Thermo Fisher, 65- 0865-14) and antibodies for 30min at 4°C: anti-CD11b (1:50, Milteny, 130-113-806), anti-hCD45 (1:50, BD Bioscience, 555485), anti-mCD45 (1:200, BD Bioscience, 563890). Samples were run on the BD FACSCanto^TM^ flow cytometer or Miltenyi MACS Quant Tyto cell sorter. After gating for CD11b^+^ cells, mouse microglia were defined as mCD45^+^ and hMG as hCD45^+^ cells. Data was analyzed using FlowJo software. Xenotransplanted hMG used for RNA sequencing were sorted on the Miltenyi MACS Quant Tyto cell sorter.

### RNA isolation

Sorted hMG were lysed in 300ul RLT buffer as supplied in the Qiagen RNeasy Kit and further processed for RNA isolation using the RNeasy Mini Kit (Qiagen 74104) following manufacturer’s instructions. Lysates were mixed with an equal volume of 70% ethanol and transferred to columns provided in the Kit. Columns were then washed sequentially with RW1 and RPE buffer provided in the Kit. As last step, RNA was eluted in 45ul RNA-free water and stored at -80°C.

### FACS

Lymph nodes, spleens and brain were collected from mice sacrificed and perfused as above, in FACS Buffer. Single-cell suspensions from lymphoid organs were prepared by mechanical dissociation. Blood, spleens and lymph nodes were processed on a 70-μm pore-sized strainer in FACS Buffer, and cells were centrifuged for 5 min at 300 x g. Red blood cell lysis was performed by using 1x red blood cell lysis buffer (420301, Biolegend). Single-cell suspensions from brain tissue were prepared as described above.

Single cells were resuspended in FACS Buffer and incubated for 15 min on ice with FcR Blocking reagent (Miltenyi, 130-092-575). Surface staining was performed for 45 min on ice using the following antibodies; anti-CD45 (Clone 30-F11, Biolegend), anti- CD3(145-2C11, Biolegend), anti-CD19 (Clone 1D3, Biolegend), anti-NK1.1 (Clone PK136, Biolegend), anti-MHCII (Clone M5114.15.2, Biolegend), anti- CD11b (Clone M1/70, BD bioscience). Death cells have been excluded from the analysis using the eBioscience Fixable Viability Dye eFluor780 (1:1000, Thermo Fisher, 65-0865-14). Cells were washed and resuspended in FACS buffer before flow cytometric analysis on a Fortessa X-20, using the FACSDiva software. Data were analyzed using FlowJo (TreeStar, Version 10.7).

### Immunohistochemistry and microscopy

The 4% PFA post-fixed brain hemispheres of hCSF1^Bdes^, *App^Hu^*and both *App^SAA^* mice were sectioned coronally on a vibratome (30µm). Sections used for Aβ staining, underwent antigen retrieval using 10mM Sodium Citrate buffer pH 6.0 (Sigma Aldrich). Sections were washed 3x 5min in phosphate-buffered saline (PBS), followed by 15min permeabilization using PBS with 0.2% Triton X-100 (PBST) and a 20min incubation at room temperature (RT) with X34 staining solution [10 µM X34 (SML1954-5MG, Sigma-Aldrich) diluted in 60% PBS (vol/vol), 40% ethanol (vol/vol) and 20mM NaOH (Sigma-Aldrich)]. Following 3x 2min washes using 60% PBS (vol/vol) with 40% ethanol (vol/vol), and 2x 5min PBST washes, sections for HLA/CD9 staining were incubated with 3% H2O2 in PBS for 1h to quench endogenous peroxidases, followed by blocking and permeabilization of all sections in blocking buffer (5% donkey serum in PBST) for 1h at RT. After overnight incubation with the primary antibodies [1/200 anti-HLA (ab7856 Abcam), 1/1000 anti-P2RY12 (HPA014518 Atlas Antibodies), 1/200 82E1 (10323 IBL-America), 1/500 anti-iba1 (234004 Synaptic Systems), 1/200 anti-Nuclei (MAB1281 Milipore)] in blocking buffer at 4°C, sections were washed 3x 5min in PBS. Next, sections were incubated for 2h at RT in secondary antibodies in blocking buffer [donkey anti-rabbit Alexa488 (A21206, Invitrogen), donkey anti-mouse Alexa594 (A21203, Invitrogen), donkey anti-guinea pig Cy5 (706-175-148, Jackson ImmunoResearch Inc.), donkey anti-mouse Alexa647 (A31571, Invitrogen), goat anti-guinea pig Alexa488 (A11073, Invitrogen), donkey anti-rabbit Alexa555 (A31572, Fisher Invitrogen) all 1/500] and washed 3x 5min in PBS. Sections for the Aβ and hMG staining were mounted in Glycergel (C056330-2, Agilent) or FluorSave (345789, Millipore). Sections for the HLA/CD9 staining were again blocked for 1h in blocking buffer, followed by an overnight incubation with the second set of primary antibodies [1/100 anti-CD9-biotin (312112, Biolegend) and anti-P2RY12)]. The next day, sections were washed 3x 5min in PBS and incubated for 2h at RT with the secondary antibodies [1/500 donkey anti-rabbit Alexa594 (A21207, Invitrogen) and 1/500 Streptavidin Alexa488 (S32354, Thermo Scientific)], again washed 3x 5min in PBS and mounted using Glycergel. Slides were kept at 4°C until imaging. Confocal images were acquired using a Nikon AX inverted microscope driven by NIS (v5.42.06) software. For excitation, 405nm, 488nm, 561nm and 640nm laser lines were used. Images were acquired using a 4x (NA=) or 20x (NA=) objective lens and processed using Fiji/ImageJ software. Images shown in Figure.2 were acquired using a Nikon Eclipse slide scanner. Images were acquired using a 10x objective lens and processed using QuPath software.

### Western blot analysis

Fifty micrograms of cleared protein brain lysate (in 250 mM sucrose, 1 mM EGTA, 5 mM Tris–HCl pH 7.4) supplemented with 1% TX-100 and cOmplete protease inhibitor cocktail (Roche) was loaded in reducing and denaturing conditions on NuPAGE (Thermo Fisher Scientific) gels and subjected to electrophoresis. Following separation, proteins were transferred to a nitrocellulose membrane for Western blotting. Membranes were stained with Ponceau-S, imaged and blocked with 5% non-fat milk Tris-buffered saline, containing 0.1% Tween 20, and incubated with Anti-M-CSF antibody [EPR20948] (Abcam) recognizing both human and mouse CSF1, washed, and incubated with horseradish peroxidase–conjugated secondary antibodies (Bio-Rad). Blots were developed using the ECL Renaissance kit (PerkinElmer) using a LAS-3000 Imaging System From Fuji.

### CSF1 ELISA

Blood samples were collected in EDTA coated tubes and after clearing, plasma was subjected to ELISA using commercially available human CSF-1 (Thermo Fisher Scientific, EHCSF1) and mouse CSF-1 (Peprotech, 900-K245) ELISA kits according to the manufacturer’s instructions.

### Bulk sequencing

RNA concentration and purity were determined spectrophotometrically using the Nanodrop ND-8000 (Nanodrop Technologies, Wilmington, DE, USA) and RNA integrity was assessed using a Bioanalyzer 2100 (Agilent Technologies, Inc. Santa Clara, CA, USA). Per sample, an amount of 1ng of total RNA was used as input for the SMART-Seq HT PLUS protocol to generate full-length cDNA (version “011921”; Takara Bio USA, Inc) using a one-step RT-PCR, combining the reverse transcription and cDNA amplification into 1 step. Subsequently, 4ng of purified double-stranded cDNA was enzymatically fragmented and stem-loop adapters were ligated. Libraries were then amplified and indexed, generating Illumina-compatible libraries with unique dual indexes (UDIs) according to the manufacturer’s protocol (Bittencourt SA, 2010).

Sequence-libraries of each sample were finally equimolarly pooled and sequenced on AVITI 2x75 Cloudbreak High, single read (101-8-8-0) at the VIB Nucleomics Core (www.nucleomics.be). Reads were aligned against a combined *mus musculus* (mm10) and *homo sapiens* (GRCh38) database using STAR (v.2.7.11a, (Dobin et al., 2013), quality checked using FastQC (v.0.11.9, (Bittencourt SA, 2010)) and counted using featureCounts (v.2.0.1, (Liao et al., 2014). MRNAs with fewer than 10 counts in at least 5 samples were discarded, leaving 15504 human mRNAs for differential expression (DE) analysis using DESeq2 (v.1.42.1) in R (v.4.3.2.). To adjust for multiple testing, Benjamini-Hochberg p-value adjustment was performed.

### Enrichment analysis

Gene Ontology enrichment analysis was performed using DAVID (Huang et al., 2008). Enrichment for top marker gene sets for HLA (n=91 genes), DAM (n=89) and CRM (n=91) as published (Mancuso et al., 2024) was performed manually using the Chi-square test, testing for the overlap between the different datasets and the genes significantly differentially expressed after p-value adjustment (n=214 total; overlap with DAM (n=17), HLA (n=12) and CRM(n=13)), while controlling for the total observed genes in the dataset (n=15504). P-value adjustment on the chi-square obtained p-values was performed using Bonferroni.

### Statistical analysis

Exemplar histological images and flow cytometry plots were selected to closely resemble expression patterns seen overall in the experimental group. Comparisons between two groups were performed using unpaired Mann Whitney U tests. The value of n reported within figure legends represents the number of animals, unless otherwise specified. Values are represented as the mean ± SD, with differences considered significant when p < 0.05. Graphs were prepared with GraphPad Prism (GraphPad Software v9.5.1) or R (v.4.3.2).

## Results

### Generation of the mice

The successful engraftment of human microglia depends on the expression of humanized CSF1 as human CSF1R signalling cannot be fully activated by murine CSF1. To avoid rejection, depletion of T and B cells is also necessary (Shinkai et al., 1992). We used CRISPR-Cas9 to interchange the part of mouse *Csf1* gene encoding amino acids 33-552 with the human counterpart corresponding to amino acids 33-554. Regulatory elements such as the promotor, 5′UTR, murine signal peptide (amino acid 1- 32) and 3’UTR of the mouse were preserved by this targeting (Additional Figure 1A). The *Rag2* gene was targeted by a CRISPR-Cas9 strategy resulting in the deletion of amino acids 16-430 and the introduction of a premature stop codon (Additional Figure 1B). Founder mice were generated in the C57Bl6J background and backcrossed over two generations to cross out potential unwanted off target events from the gene editing. Then they were intercrossed to generate the new immune deficient mouse model that we named hCSF1^Bdes^. The mouse was crossed with *App^Hu^* and *App^SAA^* mice (Serneels et al., 2020; Xia et al., 2022). While CRISPR-Cas9 technology to generate this model is patented, most academic institutions have the required permissions that enable the use of this model. Commercial entities must ensure they have the appropriate licenses for CRISPR/Cas9 technology generated materials.

### Physiological expression levels of human CSF1 in the knock-in model

The new CSF1^Bdes^ mice are fertile, healthy, produce normal litter size and show no obvious abnormal behaviour. As physiological expression levels of CSF1 are of major importance for health (Harris et al., 2012; Kwon et al., 2022) we evaluated the expression levels of human CSF1 in brain and blood from the humanized hCSF1^Bdes^ KI mice and compared those levels with WT mice. Brain CSF1 levels analysed by western blot (Figure 1A) are comparable between the two strains. Next, we measured the levels of CSF1 in the blood from WT, homozygous and heterozygous hCSF1^Bdes^ mice by ELISA. As expected, WT mice expressed only mouse CSF1, CSF1^Bdes^ heterozygous mice expressed both mouse and human CSF1 and homozygous CSF1^Bdes^ mice expressed only human CSF1. For the measurements we used kits from two different companies with their own calibrators, which makes a direct comparison of absolute values between the measurements relative, yet values are within the reported range. Summarized, we demonstrated normal physiological expression of human CSF1, i.e. not exceeding the levels of mouse CSF1, in brain and blood.

### The novel hCSF1^Bdes^ mice lack mature lymphocytes

The hCSF1^Bdes^ mice lack the *Rag2* gene, whose gene product is an essential component of the V(D)J recombinase system which results in the absence of mature B cells and T cells. Consequently, the weights of the spleen and thymus were significantly lower in the hCSF1^Bdes^ mice than in WT mice, while body weight was unaffected (Figure 1C) resembling other *Rag2* KO strains (Roh et al., 2023; Shinkai et al., 1992) (Suppl Figure 2). The presence of live, CD45^high^ lymphoid cells was analyzed in the spleen (Figure 1 C), and in lymph nodes (Figure 1D) using flow cytometry. Expression of specific markers allowed identification of different subsets such as T cells (CD3^pos^,N K1.1^neg^, CD19^neg^), natural killer (NK) T cells (CD3^pos^ ,NK1.1^pos^, CD19^neg^), NK cells (CD3^neg^, NK1.1^pos^, CD19^neg^) and B cells (CD3^neg^, NK1.1^neg^, CD19^neg^). As expected hCSF1^Bdes^ mice lack T cells, B cells and NK T cells in lymphoid organs. hCSF1^Bdes^ mice are not devoid of innate lymphoid cells, and contained similar absolute numbers of NK cells, although because of a niche depletion effect, NK cells represent the majority of the lymphoid cells found in the lymphoid tissues (Figure1D-F). In summary, we confirmed that the hCSF1^Bdes^ mouse model recapitulates immunological features linked to RAG2 deficiency (Roh et al., 2023; Shinkai et al., 1992)

### Efficient xenotransplantation of human microglia in the model

Utilizing our previously published protocol (Fattorelli et al., 2021) we transplanted hMG progenitors derived from the embryonic stem cell line H9 (WAe009-A, Wi-Cell) in hCSF1^Bdes^ mice at postnatal day (P)4. One- and 3-months post-transplantation the percentage of hMG was estimated by flow cytometry. One month after the transplantation the percentage of hMG, defined as hCD45^pos^ cells, out of the total microglia number (hCD45^pos^ + mouse CD45^pos^) was around 55% (Fig.2A), implying half of the microglia in the mouse brain were of human origin. At three months after xenotransplantation the number of hMG reached levels up to 68% of the total pool of microglia (Fig.2B-C). Using the same protocol, we also tested hMG derived from an induced pluripotent stem cell (iPSC), namely UKBi011- A-3, and xenotransplanted them into hCSF1^Bdes^. Similarly to H9-derived microglia, we observed around 50% hMG in the brains of engrafted mice 1 and 6 months post-transplantation (Supp.Fig.S2), which resembles graft efficiencies in hCSF1^Flv^ mice (Fattorelli et al., 2021; Mancuso et al., 2024).

**Figure 2.**
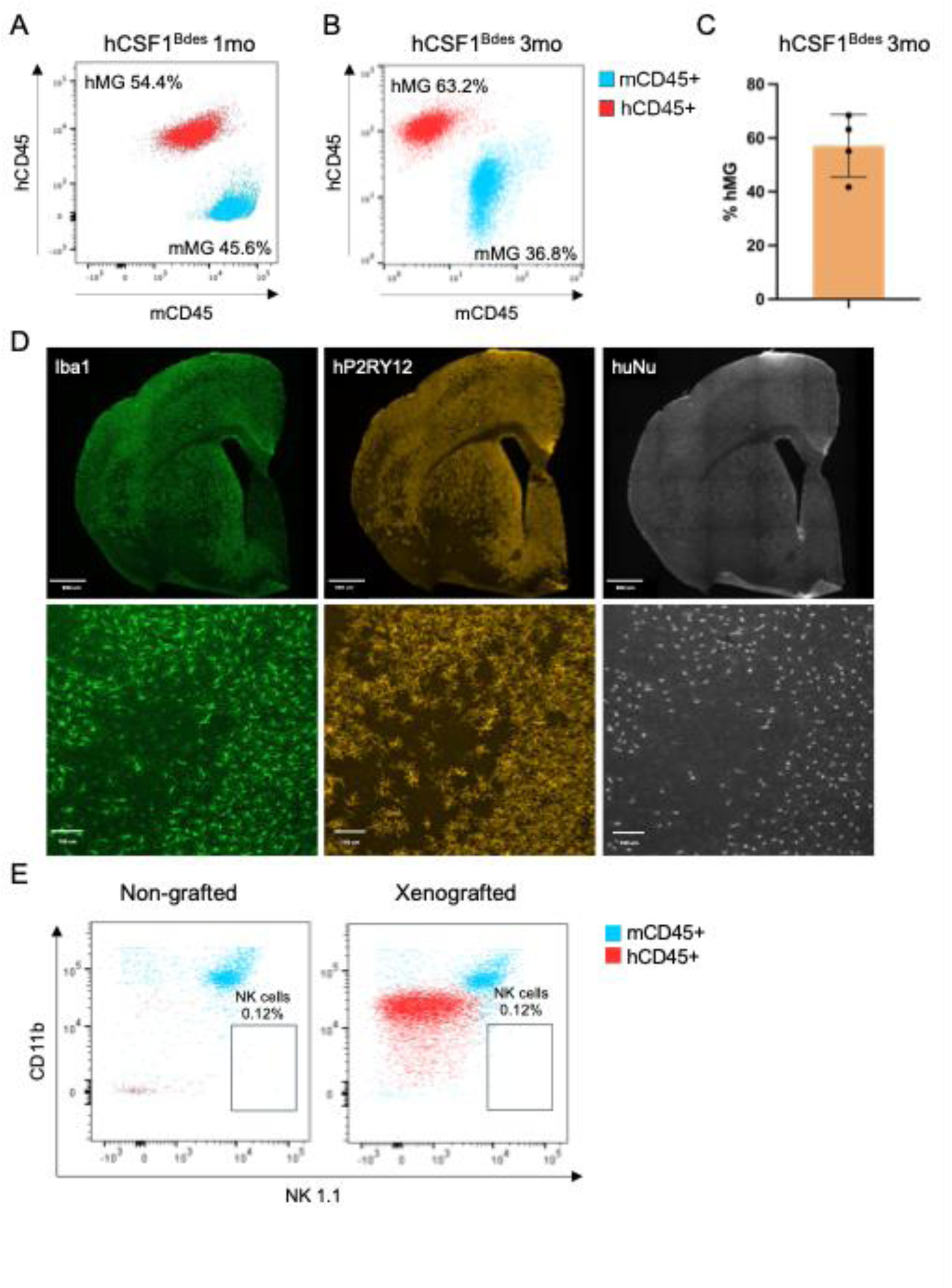
Efficient xenotransplantation of human derived microglia. A-B. hCSF1^Bdes^ mice were xenografted with human microglia. Isolated microglia were analyzed by flow cytometry one month (A) and 3 months (B) after xenotransplantation. Human microglia (hMG) are represented as hCD45+ population after gating for CD11b+ cells. Mouse microglia (mMG) are represented as mCD45+ population after gating for CD11b+ cells. Samples were acquired on BD FACSCanto flow cytometer (A) or Miltenyi Quant Tyto cell sorter (B) and analysed in FlowJo software. C. % of hMG in 3mo xenografted hCSF1^Bdes^. N=4, Bar plot represents Mean ± SD. D. Representative images of engrafted human microglia in the brain of hCSF1^Bdes^ mice at 3 months of age. Microglia cells (both mouse and human) are identified as Iba1+ cells. Xenografted human microglia are identified as hP2RY12 and huNu positive cells. Scalebar, 1mm (top) and 100µm (bottom). E. Xenografting of human microglia in hCSF1^Bdes^ mice does not induce NK cells expansion in brain, 6 months after grafting. Representative flow cytometry plots displaying absence of NK 1.1 positive cells in hCSF1^Bdes^ mice xenografted with human microglia (right), and in hCSF1^Bdes^ without grafted cells (left).

We further investigated spread and morphology of transplanted hMG by immunohistochemistry. Microglial cells, both mouse and human, were stained using the well-established Iba1 marker. HMG were further distinguished using the human-specific P2RY12 antibody, and a human nuclei (huNu) staining. We observed wide spread distribution of hMG, defined as P2RY12^pos^ and huNu^pos^ cells, in the mouse brain 3 month after transplantation (Fig.2D). Furthermore, engrafted hMG showed their typical ramified microglial morphology (Fig.2D) as described before (Hasselmann et al., 2019; Mancuso et al., 2019). We also confirmed that xenografting of hMG in hCSF1^Bdes^ mice does not induce NK cells expansion in brain (Fig.2E)

In summary, we confirm that human stem-cell derived microglial progenitors can be successfully xenotransplanted into the brains of hCSF1^Bdes^ mice. After the xenotransplantation, hMG progenitors acquire their typical ramified morphology and colonize the mouse brain efficiently.

### The new model enables to study the hMG response to Alzheimer’s Aβ pathology

We next assessed the utility of the Csf1^Bdes^ mouse for researching the hMG response to amyloid-beta pathology by crossing it to the freely available *App^SAA^* mouse (Xia et al., 2022) as well as a control mouse that carries the humanized amyloid-beta sequence without any pathological mutations (referred to as the *App^Hu^* mice (Serneels et al., 2020)). Both mouse strains were xenotransplanted with H9-derived hMG at P4 and graft efficiency in both mouse strains was assessed at 1 and 6 months post xenotransplantation, showing similar graft efficiencies for both strains (see Fig.3A-C). Additionally, staining the mouse brain for hMG using the hMG marker hP2RY12 showed wide spread distribution of hMG in 6-month-old *App^Hu^* and *App^SAA^*mice, with hMG inhabiting areas with amyloid plaque deposits in *App^SAA^*mice (Fig.3D) as defined by 82E1^pos^ and/or X34^pos^ areas.

**Figure 3.**
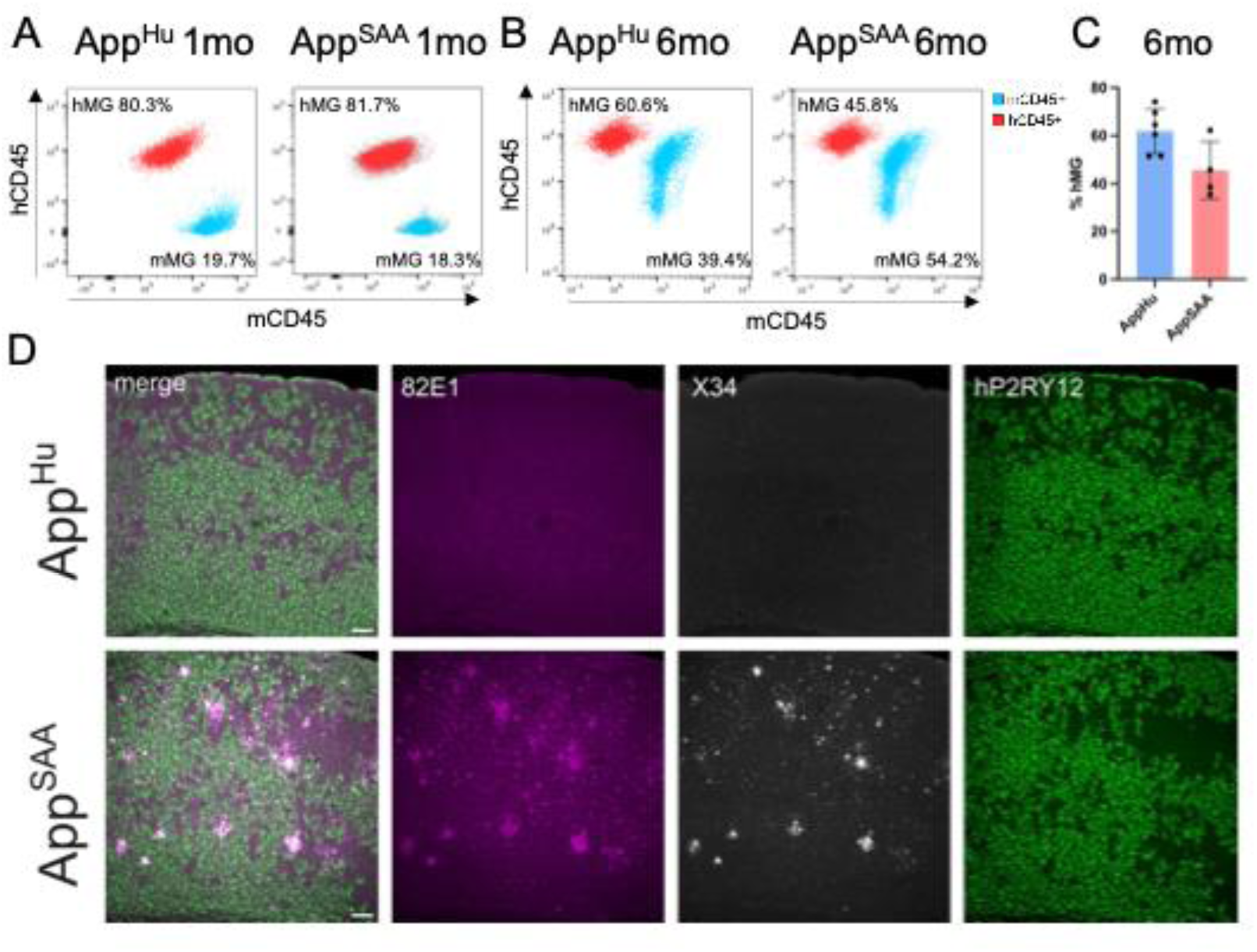
Human microglial response to Aβ pathology. A-B. Representative flow plots for xenografted *App^Hu^* (left) and *App^SAA^* (right) mice 1 month (A) and 6 months (B) after human microglia transplantation. Human microglia (hMG) are represented as hCD45+ population after gating for CD11b+ cells. Mouse microglia (mMG) are represented as mCD45mid cells after gating for CD11b+ cells. Samples were analysed on BD FACSCanto flow cytometer (A) or Miltenyi Quant Tyto cell sorter (B) and analysed in FlowJo software C. % of hMG in 6mo xenografted App^Hu^ and App^SAA^. N=4-6, Bar plot represents Mean ± SD. D. Representative images of human microglia engrafted in the cortex of App^Hu^ and App^SAA^ mice at 6 months of age, labeled with human-specific antibodies for hP2RY12, as well as X34 and 82E1 for Aβ plaques. Scale bar, 100µm.

At 6 months of age, the response of hMG to amyloid pathology was assessed by bulk RNA sequencing on sorted hMG and immunohistochemistry. Differential expression analysis comparing the transcriptome of sorted hMG xenotransplanted in the *App^Hu^* vs the *App^SAA^* mice revealed 214 significantly differentially expressed genes (padj <0.05), with 152 upregulated and 62 downregulated genes. Among the top upregulated genes were *CD83*, *RGS16* and *CD9* whereas *CLDN4, TMEM176B* and *SOX5* were most downregulated (Fig.4A).

**Figure 4.**
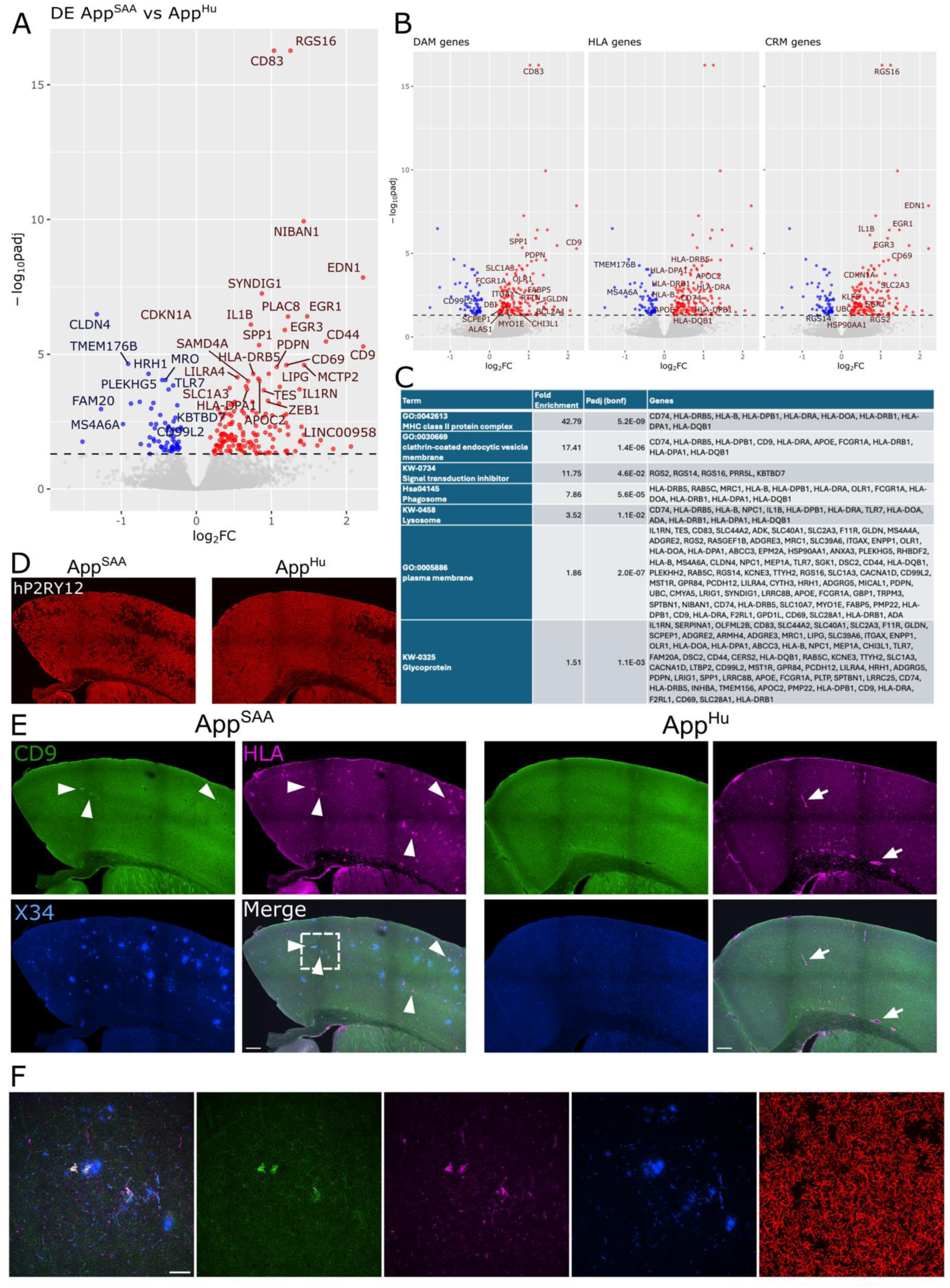
Bulk RNAseq demonstrates the activation of huMG in response to Aβ pathology. A. Vulcano plot of the most differentially up- (red) and down-regulated (blue) genes (Benjamini- Hochberg corrected p-value (padj)<0.05) when comparing the bulk RNAseq profile of sorted huMG engrafted in *App^SAA^* vs *App^Hu^* mice at 6 months of age. B. Vulcano plots highlighting the top 100 marker genes that were significantly differentially expressed (padj<0.05) in this dataset for disease- associated microglia (DAM), antigen presenting microglia (HLA) and cytokine response microglia (CRM1) and as found in (Mancuso et al., 2024). C. Significantly enriched gene ontology (GO) categories, Uniprot Keywords (UP-KW) and KEGG pathways as found by DAVID. D-F. Immunohistochemistry for hP2RY12 (D), hCD9, hHLA (HLA-DR/DP/DQ) and X34 (E-F) in 6 month old App^SAA^ vs App^Hu^ mice. D-E; scalebar visible in merge: 200 µm. E. arrow-heads (left panel) pinpoint parenchymal plaque-associated hHLA and hCD9 staining in App^SAA^ mice, whereas arrows (right panel) indicate blood-vessel associated staining for hHLA in App^Hu^ mice. F. Zoom of insert in merge from panel E, scale bar: 50 µm.

Our lab has demonstrated previously that the hMG response to Aβ pathology is more complex and multi-faceted than the mouse response (Mancuso et al., 2024). By using the top marker genes for disease-associated microglia (DAM), antigen presenting microglia (HLA) and cytokine response microglia (CRM) from Mancuso et al. (Mancuso et al., 2024) we could demonstrate a significant enrichment of DAM (Chi-square (1, n=15504)=147.6, padj=6.6e-16), HLA (Chi-square (1, n=15504)=69.8, padj=6.6e-16) and CRM genes (Chi-square (1, n=15504)=82.9, padj=6.6e-16) among the DE genes, demonstrating the ability of hMG within the *App^SAA^* mice to mount a transcriptional response to amyloidosis that is similar to what we described previously in the *App^NLGF^* mice (Fig.4B). GO enrichment analysis using DAVID (Fig.4C) further confirms that hMG within the *App^SAA^*mice deregulate genes associated with the MHC-II response, clathrin-mediated endocytosis, phagosome and lysosome, which is in line with previously reported differentially expressed genes in hMG when facing amyloid pathology (Claes et al., 2022; Mancuso et al., 2024). By using immunohistochemistry we again confirm similar graft efficiencies between the strains (Fig.4D) and find that 6 month old *App^SAA^* mice display robust plaque pathology in regions xenotransplanted with hMG. We found that whereas expression of CD9 and MHC-II genes was mainly restricted to the blood vessels in *App^Hu^* mice (white arrows Fig.4E right panel), *App^SAA^* mice display an upregulation of CD9 and MHC-II protein in the brain parenchyma and particularly in close vicinity to plaques (white arrow heads Fig.4E left panel & zoom Fig.4F), which is in line with our bulk RNAseq data and previous reports using other models of amyloidosis (Claes et al., 2021, 2022; Hasselmann et al., 2019; Mancuso et al., 2024). In conclusion, these results demonstrate that the new Csf1^Bdes^ mouse provides a valuable alternative for xenotransplantation experiments and studying the response of hMG to AD-relevant pathologies.

## Discussion

Microglia xenotransplantation can be performed with success in the hCSF1^Flv^ mice (Rag2tm^1.1Flv^; Csf1^tm1(CSF1)Flv^;Il2rg^tm1.1Flv/J^) (Hasselmann et al., 2019; Mancuso et al., 2019). Humanization of CSF1 is required for the integration, survival and proliferation of human myeloid cells in the mouse environment (Abud et al., 2017; Fattorelli et al., 2021; Hasselmann et al., 2019; Mancuso et al., 2019, 2024; Svoboda et al., 2019; Xu et al., 2020). Conversely, deletion of both *Rag2* and *Il2rg* genes ensures the absence of functional T cells, B cells and NK cells, which makes the Rag2tm^1.1Flv^;Il2rg^tm1.1Flv/J^ the elective model for xenotransplantation. The hCSF1^Bdes^ mouse model that we generated expresses the *hCSF1* and lacks the *Rag2* gene, and therefore it lacks functional T cells and B cells, while NK are present in normal numbers (Figure 1). In contrast to T and B cells, NK cells represent the innate lineage of lymphocytes that possess germline-encoded antigen receptors and they do not require RAG proteins for their development, function, or survival (Kondo et al., 1997; Shinkai et al., 1992). Early activation of NK cells following transplantation is associated with killing of allogeneic target cells and release of immunomodulatory chemokines and cytokines, which can contribute to rejection. However, there is no evidence that IL2ry deficiency is needed for human cell transplantation in brain (Benichou et al., 2011).

We demonstrate here that we can successfully transplant iPSC-derived hMG into the brain of hCSF1^Bdes^ mice (Figure 2), without eliciting an NK driven immune response. hMG could survive and were able to stably colonize a large area of the mouse brain (>50% of all microglia were human), and showed the typical ramified morphology and expression of homeostatic markers comparable to similar experiments performed in the hCSF1^Flv^ mice (Mancuso et al., 2019, 2024; Xu et al., 2020a). We further show that by crossing the Csf1^Bdes^ mouse to the *App^SAA^* mouse model of AD and *App^Hu^* control mice, the xenografted hMG exposed to amyloid-beta pathology in *App^SAA^* upregulate the typical activation markers previously described, including CD9 and HLA-DQ/DR both on mRNA and protein levels, predominantly in the vicinity to plaques. Our data largely agree with our previous work confirming that the described phenotypes of the multipronged amyloid plaque response are maintained in this independent novel mouse model for studying human microglia responses in disease (Mancuso et al., 2024).

Thus, our Csf1^Bdes^ mouse proves to be a valuable new alternative for the broader research community. Whereas the current standard in the field, the hCSF1^Fvl^ is restricted to non-profit entities as dictated by the Regeneron patent EP2675271B1, our model demonstrates that the combination with Il2rg^tm1.1Flv/J^ is not needed for microglia transplantation. We furthermore humanized the CSF1 locus and generated an entirely novel Rag2 targeted allele. Users should take into account that Crispr/Cas9 technology was used to generated this novel model, and investigators need to check whether their institute is subject to the terms and conditions of the limited license from the Broad institute and from Caribou Biosciences, Inc. Thus, in principle, the Csf1^Bdes^ mouse can be used by non-profit and for-profit organisations, facilitating the use of the xenotransplantation paradigm for hMG to study complex human disease. We also anticipate that this will further enable the exploration of the use of hMG transplantation as a therapeutic modality for various diseases (Chadarevian et al., 2024; Munro et al., 2024) Particularly in combination with the newly developed CSF1R inhibitor-resistant human CSF1R variant (Chadarevian et al., 2023)) and the broader advances the scientific community is making on the use of transplantation models for the treatment of neurological disorders such as epilepsy or Parkinson disease(Svendsen 2024 Nat Med), stem cell-based therapies using microglia may hold a place in disease treatment in the not-too distant future.

## Conclusion

Our hCSF1^Bdes^ model demonstrates that NK cell depletion is not required for successful human microglia transplantation and further work validated the multipronged response of human microglia to amyloid plaques. This versatile new model is accessible to both non-profit and for-profit organizations, facilitating the broader application of the xenotransplantation paradigm with human microglia (hMG) to advance research on complex human diseases.

## List of abbreviation

AD: Alzheimer disease
CSF1: Colony stimulating factor 1
hMG: human Microglia
IL2RG: Interleukin 2 Receptor Subunit Gamma
NK: Natural killer
RAG: recombination activating gene
SD: standard deviation

## Declarations Ethics approval

All animal experiments were conducted according to protocols approved by the local Ethical Committee of Laboratory Animals of KU Leuven (P125-2022). The use of human ESC (H9) and iPSC (UKBi011-A-3) are approved by the local Ethical Committee of UZ Leuven (S63481).

## Consent for publication

Not applicable

## Availability of data and materials

Bulk sequencing data are deposited in the Gene Expression Omnibus (accession number) and from the B.D.S lab website (URL) . Mice are deposited to the European Mutant Archive (EMMA). All materials used in the current work can be obtained upon request to B.D.S. and will be made available under the standard MTA of VIB.

## Authors’ contributions

L.S., A.Si., E.P., I.G. and B.D.S conceived and designed the study and wrote the manuscript. L.S., A.Si., E.P., I.G., A.M.M, A.N., A.Sn. performed experiments and analyzed data together with B.D.S. All authors discussed the results and commented on the manuscript.

## Acknowledgement

The authors thank Amber Claes and Veronique Hendricks for breeding and taking care of the mice. The VIB-KU Leuven Center for Brain & Disease Research Technologies, Mouse Expertise Unit for rederivation of the mouse colonies. This work was funded by a FBRI research funding awarded to B.D.S. entitled “Generation and initial characterization of a humanized CSF1 Knock in model to study microglia function and dysfunction”. This project received further funding from the European Research Council (ERC) under the European Union’s Horizon 2020 Research and Innovation Programme (grant agreement no. ERC-834682 CELLPHASE_AD). This work was also supported by funding from UKRI and the Medical Research Council (MR/Y014847/1) via the Dementia Research Institute. Further support was given by the Flanders Institute for Biotechnology (VIB vzw), a Methusalem grant from KU Leuven and the Flemish Government, the Fonds voor Wetenschappelijk Onderzoek, KU Leuven, The Queen Elisabeth Medical Foundation for Neurosciences, the Opening the Future campaign of the Leuven Universitair Fonds, The Belgian Alzheimer Research Foundation (SAO-FRA) and the Alzheimer’s Association USA. B.D.S. holds the Bax-Vanluffelen Chair for Alzheimer’s Disease.

