## Additional figures for "A versatile mouse model to advance human microglia transplantation research in neurodegenerative diseases"

### Additional Data

*Equal contributions

#### Additional figure 1

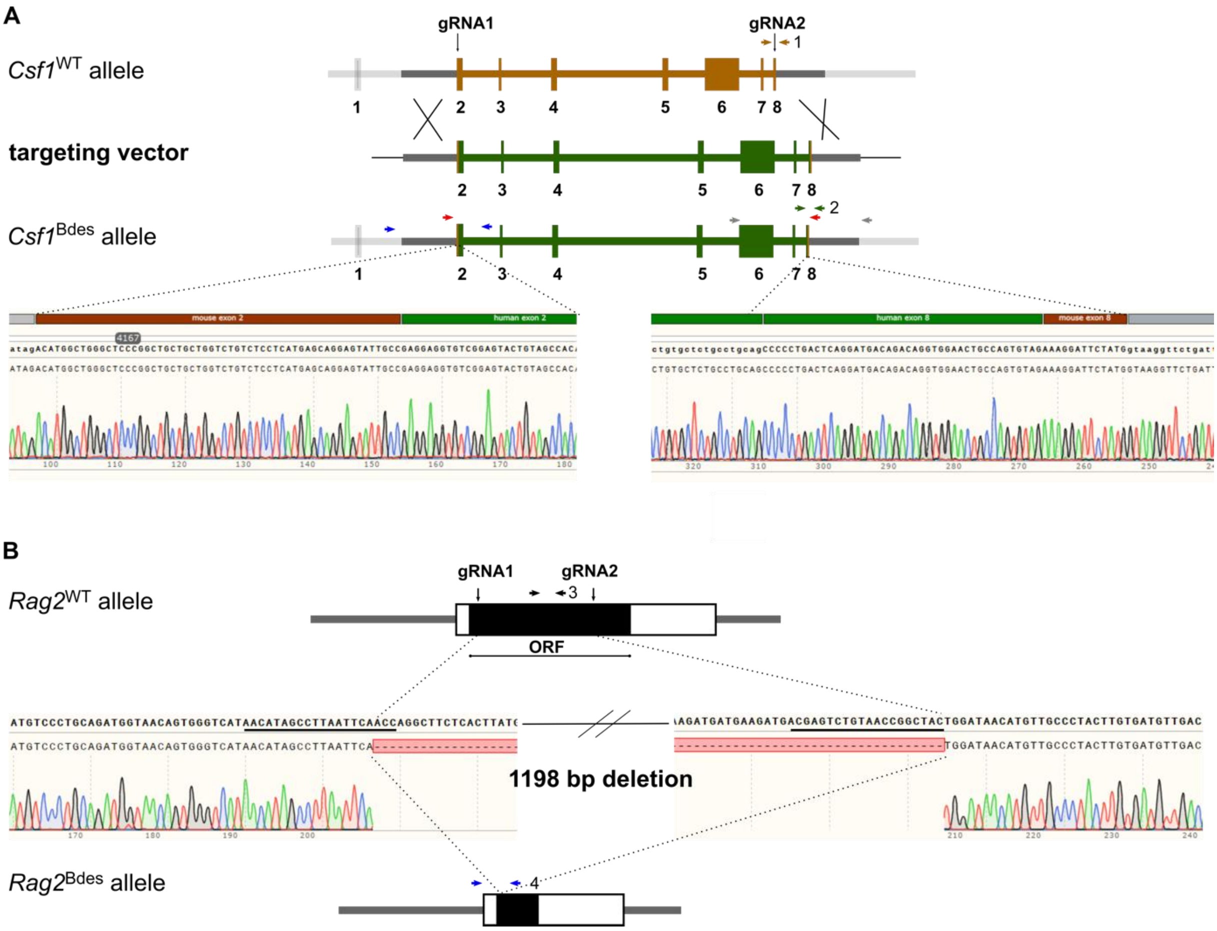


**Additional Figure 1** Generation of the humanized Csf1 and Rag2 knock mice

A. Schematic overview of the strategy used to generate the humanized *Csf1* KI allele. Exons are presented as boxes; The positions of the guide RNA's are indicated. Primers used for quality control by PCR and Sanger sequencing are depicted as arrows. The mouse exons and introns presented in brown are replaced by human exons and introns indicated in green. The lower panel shows the Sanger sequencing results at the recombination sites. Genotyping primers are indicated with brown arrows (1) for the *Csf1*$^{WT}$ allele and green (2) for the *Csf1*$^{Bdes}$ allele. B. Schematic overview of the strategy used to generate the *Rag2*$^{Bdes}$ KO allele. Exon3 of the mouse *Rag2* gene is depicted as a box, the complete open reading frame of the Rag2 gene as a black box, the position of the two guide RNA's are indicated. The lower part shows the Sanger sequencing results at the deletion site demonstrating that 1198 bp of the coding sequence are deleted. The primers used to genotype the alleles are indicated with arrows black arrows (3) for *Rag2*$^{WT}$ allele and blue arrows (4) for *Rag2*$^{Bdes}$ allele.

**Additional figure 1**

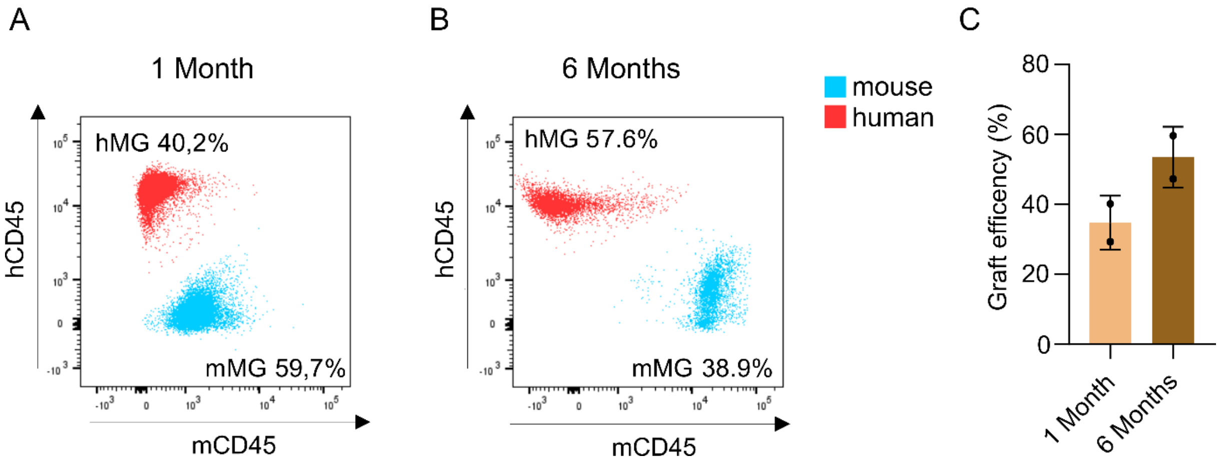


**Additional Figure 2** Efficient xenotransplantation of human derived microglia in the hCSF1$^{Bdes}$ mouse model.

A-B. hCSF1$^{Bdes}$ mice were xenografted with human microglia derived from iPSCs ( UKBi011-A-3). Isolated microglia were analyzed by flow cytometry one month (A) and 6 months (B) after xenotransplantation. Human microglia (hMG) and mouse microglia (mMG) are represented as percentage of total CD11b+ cells. C. Graft efficiency of hMG at 1 and 6 months after transplantation. N=2, Bar plot represents mean ± SD.
